# TCGA Pan-Cancer genomic analysis of Alternative Lengthening of Telomeres (ALT) related genes

**DOI:** 10.1101/2020.04.27.063610

**Authors:** Isaac Armendáriz-Castillo, Andrés López-Cortés, Jennyfer García-Cárdenas, Patricia Guevara-Ramírez, Paola E. Leone, Andy Pérez-Villa, Verónica Yumiceba, Ana K. Zambrano, Santiago Guerrero, César Paz-y-Miño

## Abstract

Telomere maintenance mechanisms (TMM) are used by cancer cells to avoid apoptosis, 85-90% reactivate telomerase, while 10-15% use the alternative lengthening of telomeres (ALT). Due to anti-telomerase-based treatments, some tumors have the ability to switch from a telomerase-dependent mechanism to ALT, in fact, the co-existence between telomerase and the ALT pathway have been observed in a variety of cancer types. Despite different elements in the ALT pathway have been uncovered, the molecular mechanism and other factors are still poorly understood, which difficult the detection and treatment of ALT-positive cells, which are known to present poor prognosis. Therefore, with the aim to identify potential molecular markers to be used in the study of ALT, we combined simplistic *in silico* approaches in 411 telomere maintenance (TM) genes which have been previously validated or predicted to be involved in the ALT pathway. In consequence, we conducted a genomic analysis of these genes in 31 Pan-Cancer Atlas studies (n=9,282) from The Cancer Genome Atlas in the cBioPortal and found 325,936 genomic alterations, being mRNA high and low the top alterations with 65,.8% and 10.7% respectively. Moreover, we analyzed the highest frequency means of genomics alterations, identified and proposed 20 genes, which are highly mutated and up and down regulated in the cancer studies and could be used for future analysis in the study of ALT. Finally, we made a protein-protein interaction network and enrichment analysis to obtain an insight into the main pathways these genes are involved. We could observe their role in main processes related to the ALT mechanism like homologous recombination, homology directed repair (HDR), HDR through homologous recombination and telomere maintenance and organization.. Overall, due to the lack of understanding of the molecular mechanisms and detection of ALT-positive cancers, we identified and proposed more molecular targets that can be used for expression analysis and additional *ex vivo* assays to validate them as new potential therapeutic markers in the study of the ALT mechanism.

## Introduction

Telomeres are nucleoprotein complexes that consist of a tandem 5’-TTAGGG-3’ sequence and protect the ends of eukaryotic chromosomes preventing DNA damage response (DDR), end-to-end fusions and genomic instability ^1^. Telomeric DNA ranges from 3 to 15 Kb in humans, leaving a 3’-single-strand overhang, usually called the G-overhang ^1,2^. To avoid this end-replication problem, telomeres are protected by a complex of proteins called shelterin, which is essential in the formation of the t-loop and hides the G-overhang ^2^. However, with each round of somatic cells cycle, telomeres lose about 200 nucleotides; eventually, this shortening leads to senescence or apoptosis ^3^.

To avoid apoptosis or senescence due to telomere shortening, cancer cells use a set of mechanisms known as Telomere Maintenance Mechanisms (TMM), which includes: telomerase reactivation and the Alternative Lengthening of Telomeres (ALT) ^1,3^. A high proportion of tumors reactivate the expression of telomerase to maintain its chromosomal ends, however, 10-15% of human cancers use the ALT pathway ^4^.

ALT-positive (ALT+) cells display a common characteristic phenotype. For instance, telomeric DNA in ALT+ cells is constantly elongating, showing that its telomeres rely on recombining mechanisms like homologous recombination (HR) and homology-directed repair (HDR) pathways to be extended ^5,6^. ALT+ cells also show heterogenous telomere length, abundant extrachromosomal repeats (ECTRs), telomere sister chromatid exchange (T-SCE) ^7^ and high levels of extrachromosomal telomeric single-stranded DNA known as C-circles, which are markers for ALT+ cells detection ^8^.

The main characteristic of ALT+ telomeres is its association with promyelocytic leukemia (PML) proteins, which altogether are believed to function as platforms for telomere recombination and are known as ALT-associated PML bodies (APBs) ^7,9^. In fact, it has been shown that disruption of APBs blocks the ALT mechanism ^7^.

Although, many proteins have been implicated in the ALT mechanism, like the loss of alpha thalassemia/mental retardation syndrome X-linked chromatin remodeler (ATRX), the loss of histone chaperone death domain-associated protein (DAXX) and histone H3.3 (H3F3A) and the well-known expression of RAD51/52 complex, the molecular basis through which ALT occurs remain elusive and poorly understood ^1,5,10^.

Many anti-telomerase-based cancer therapies used in cancer treatment, are believed to cause the switching in some tumors from telomerase to ALT ^3,11^. Indeed, the co-existence of telomerase and the ALT mechanism has been reported in some cancer types ^3,12^. This switching has converted ALT in a potential target for therapy in the last years, due to the poor prognosis these cells represent ^1,7,13^. Different drugs have been tested in the last years to target ALT+ cells, however, most of them have been unsuccessful or are on phase 1/2 of clinical trials^1,7^.

Although several ALT-associated factors have been uncovered, the pathway activation, regulators and functioning require further investigation ^7^. Therefore, the identification of new potential molecular markers would be of enormous importance in the future design of new strategies for the detection and treatment of ALT+ cancers. To fulfill this need, we evaluated the genomic alterations of 411 telomere maintenance (TM) genes in 9,282 samples from 31 Pan-Cancer Atlas (PCA) studies and applied different *in silico* approaches with the aim to identify and propose more genes which after validation could be used as potential molecular markers to improve the undersantinding of the ALT pathway.

## Results

### Genomic alterations

To address the genomic alterations of TM genes, a total of 411 genes from the TelNet database^14^ (Table S1) were analyzed in the cBioPortal^15,16^ by selecting 9,282 samples from 31 Pan-Cancer Atlas studies from The Cancer Genome Atlas (TCGA)^17–26^ (Table 1).

**Table 1.** List of TCGA Pan-Cancer Studies with the number of individuals.

| TCGA Study | n | TCGA Study | n |
| --- | --- | --- | --- |
| Acute Myeloid Leukemia (LAML) | 165 | Lung Squamous Cell Carcinoma (LUSC) | 466 |
| Adrenocortical Carcinoma (ACC) | 76 | Mesothelioma (MESO) | 82 |
| Bladder Urothelial Carcinoma (BLCA) | 402 | Ovarian Serous Cystadenocarcinoma (OV) | 201 |
| Brain Lower Grade Glioma (LGG) | 507 | Pancreatic Adenocarcinoma (PAAD) | 168 |
| Breast Invasive Carcinoma (BRCA) | 994 | Pheocromocytoma and Paraganlioma (PCPG) | 161 |
| Cervical Squamous Cell Carcinoma (CESC) | 275 | Prostate Adenocarcinoma (PRAD) | 488 |
| Cholangiocarcinoma (CHOL) | 36 | Sarcoma (SARC) | 251 |
| Colorectal Adenocarcinoma (COAD) | 524 | Skin Cutaneous Melanoma (SKCM) | 363 |
| Diffuse Large B-cell Lymphoma (DLBC) | 39 | Stomach Adenocarcinoma (STAD) | 407 |
| Esophageal Adenocarcinoma (ESCA) | 181 | Testicular Germ Cell Tumors (TGCT) | 144 |
| Glioblastoma Multiforme (GBM) | 145 | Thymoma (THYM) | 119 |
| Head and Neck Squamous Cell Carcinoma (HNSC) | 488 | Thyroid Carcinoma (THCA) | 480 |
| Kidney Renal Clear Cell Carcinoma (KIRC) | 352 | Uterine Carcinosarcoma (UCS) | 56 |
| Kidney Renal Papillary Cell Carcinoma (KIRP) | 274 | Uterine Corpus Endometrial Carcinoma (UCEC) | 507 |
| Liver Hepatocellular Carcinoma (LIHC) | 348 | Uveal Melanoma (UVM) | 80 |
| Lung Adenocarcinoma (LUAD) | 503 |  |  |

A total of 325,936 genomic alterations were identified (Table S2) and a pie chart with the most frequent alterations was constructed after all values were normalized by the number of samples. Figure 1A shows, mRNA upregulation at the top with 65.8%, followed by mRNA downregulation (10.7%), copy number alteration (CNA): amplifications (9.6%), missense mutation (putative passenger) with 7.2%, deep deletion (3.1%) and truncating mutations and fusion genes with less than 2%.

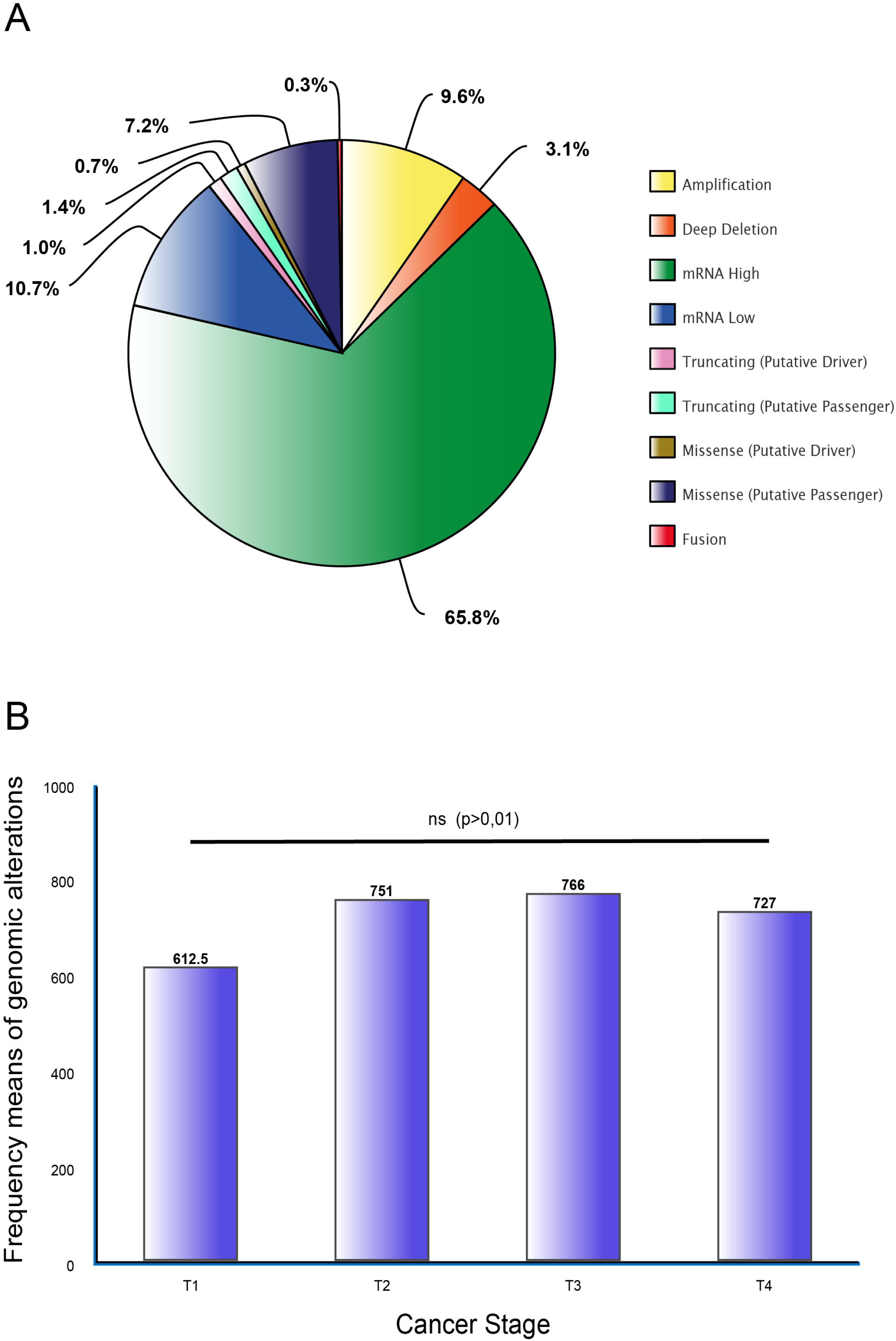

Finally, to better understand the implication of these genes in cancer progression, from primary tumors to metastasis, we analyzed staging (Figure 1B) (Table S3). However, no significant difference was observed after a Bonferroni correction test (p>0.01). As a result, it can be inferred that TM genes alterations and the ALT pathway across different types of tumors are not dependent on cancer staging.

### TM genes validation and TCGA Pan-Cancer studies frequencies

The gene set and each TCGA study were ordered by the highest frequency mean of genomic alterations to the lowest (Tables S4). To narrow down our analysis, the first quartile of genes (n=103) were selected for further analysis and the highest frequencies of alterations for the 31 PCA were: UCS (56.054), OV (53.194), UCEC (49.387), ESCA (47.088), BLCA (44.883), ACC (44.658), SKCM (44.620), LUSC (43.822), STAD (43.005), BRCA (42.068), LUAD (40.942), CESC (40.116), COAD (39.038), SARC (37.418), LIHC (37.353), DLBC (37.256), HNSC (36.381), CHOL (35.444), PAAD (33.256), MESO (30.634), UVM (30.225), GBM (29.366), TGCT (29.132), KIRP (28.708), PRAD (28.258), LGG (27.387), PCPG (26.292), KIRC (24.168), LAML (21.255), THYM (20.185) and THCA (19.319) (Figure 2A).

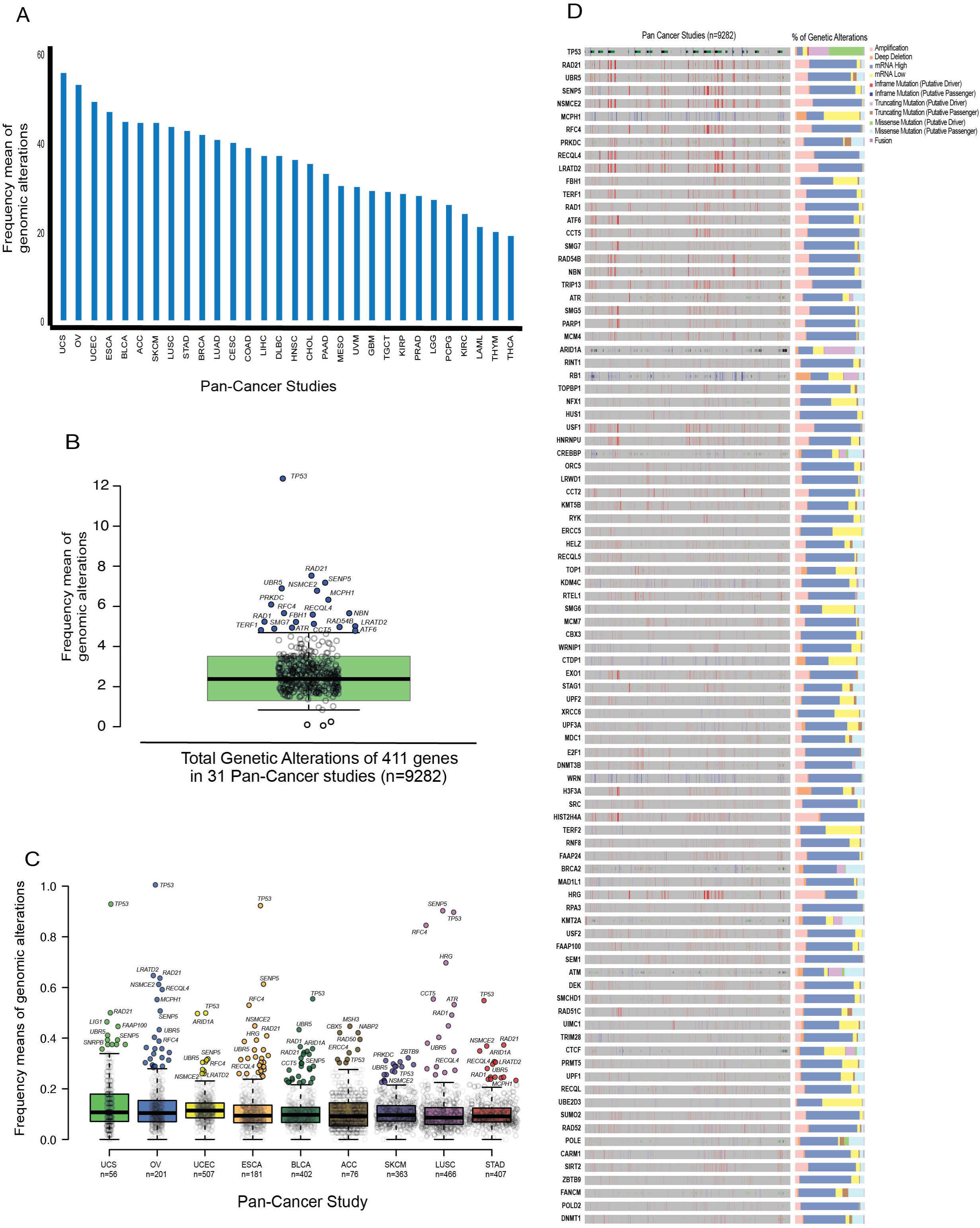

To further identify which TM genes were the most altered in the 31 PCA studies overall, a boxplot with the frequency means of genomic alterations was constructed for the 411 genes (Figure 2B). Hence, after the boxplot analysis the following 20 TM genes were selected and proposed as candidate genes to be studied in the ALT pathway: *TP53* (12.370), *RAD21* (7.529), *SENP5* (7.179), *UBR5* (6.897), *NSMCE2* (6.779), *MCPH1* (6.332), *PRKDC* (6.091), *RFC4* (5.662), *RECQL4* (5.656), *NBN* (5.587), *FBH1* (5.239), *RAD1* (5.221), *CCT5* (5.126), *LRATD2* (5.013), *RAD54B* (4.969), *ATR* (4.940), *SMG7* (4.888), *TERF1* (4.817), *ATF6* (4.778) and *ARID1A* (4.357).

Then, to obtain insights of the most altered genes in each of the PCA studies and to correlate them with Figure 2B, an additional boxplot was constructed (Figure 2C), along with the alteration frequencies of the 411 TM genes and the first quartile of the PCA studies with the highest means of genomic alterations (UCS, OV, UCEC, ESCA, BLCA, ACC, SKCM, LUSC and STAD). On this basis, the following TM genes were identified to be highly altered in more than one of the 9 types of tumors: *TP53* (9), *UBR5* (8), *RAD21* (6), *SENP5* (6), *NSMCE2* (5), *RFC4* (5), *MCPH1* (4), *RECQL4* (4), *ARID1A* (3), *CCT5* (3), *LRATD2* (3) and *RAD1* (3).

Finally, in order to elucidate the predominant genomic alteration in the first-quartile of the most altered TM genes across the 31 PCA studies, an oncoprint with the percentage of each alteration is showed in Figure 2D. As expected, mRNA upregulation is the most common alteration in the gene set, with certain exceptions where amplifications, mRNA downregulation and truncating mutations, appear most frequently.

### Protein-protein interaction (PPi) network and enrichment analysis

PPi networks are useful resources to understand how proteins interact between them in the cell ^27^, hence, STRING database ^28^ was used to observe the interactions of 103 TM proteins. By using an interaction score of highest confidence (0.9)^29^, a network was constructed, additionally, the most significant pathways (p< 0.001) were selected and marked with different colors in the nodes (Figure 3A). As a result, it became evident that 68% of the proteins in the network are essential for DNA binding, 59% for DNA metabolic process, 47% for DNA repair, 23% for telomere organization and maintenance, 12% for telomeric DNA binding and 22% are involved in DNA Double-Strand Break Repair, Homology Directed Repair (HDR) and HDR through Homologous Recombination (HRR).

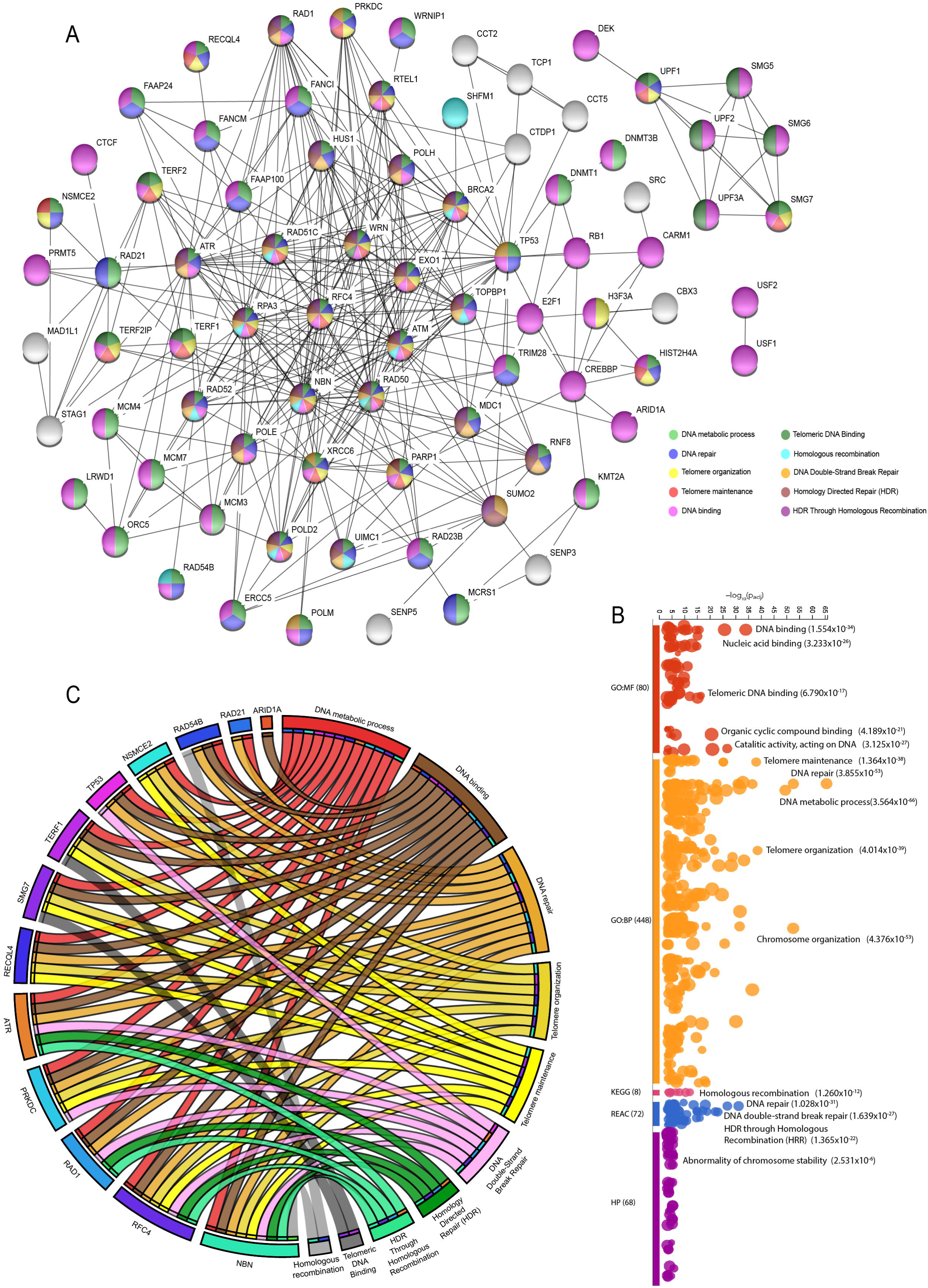

Subsequently, an enrichment analysis was made using g:Profiler^30^ (Table S5), Figure 3B shows the enrichment map of 103 TM proteins. The most significant pathways with Benjamini-Hochberg correction and False Discovery Rate (FDR) < 0.001 from Gene Ontology (GO) Molecular Function (MF) and Biological Process (BP), Kyoto Encyclopedia of Genes and Genomes (KEGG), REACTOME database and Human Phenotype (HP) are also showed^31^.

Finally, the proteins with the highest means of genomic alterations in the boxplot analysis (Figures 2B and 2C), were correlated with its most significant pathways using a CIRCOS plot (Figure 3C). As can be observed, most of these are involved in DNA repair, binding and metabolic processes, but, at the same time, it can be observed that several of the candidate proteins like *RAD54B, NSMCE2, TERF1, SMG7, RECQL4, ATR, PRKDC, RAD1, RFC4* and *NBN* which are highly altered in the PCA studies, are involved in telomere organization, maintenance and binding, HR, HDR and HRR.

Overall, the correlation of the most significant pathways observed in the different analyses where the 20 TM candidate proteins are involved, suggests a huge impact of these proteins’ alterations in the activation and progression of the ALT mechanism, which will be discussed later.

## Discussion

TM is a crucial mechanism in the hallmarks of cancer for indefinite replicative potential, 10 to 15% of cancers do not depend on telomerase to maintain or extend its telomeres, instead, they use the ALT pathway^1^. In the last years, efforts to identify genes responsible for ALT progression have been made; so far, the loss of *ATRX* and *DAXX* ^1,10,32–34^and expression of *RAD51* and *RAD52* ^1,3,4^ have been widely described. Nevertheless, ALT mechanism and the molecular basis underlying its progression, switching, detection and treatment remain elusive, therefore, we used simplistic OncoOmics and *in silico* approaches to identify potential molecular markers to improve the study of the ALT mechanism.

In 2018, TelNet database was introduced, offering more than 2000 human genes associated with TM. Genes are annotated according to TM mechanism, function, significance score and its validation in the literature^14^. We manually curated the database by selecting 411 TM genes which have been predicted or validated to be involved in the ALT pathway. So far, the main recurrent mutations identified in ALT+ cells are associated with *ATRX*/*DAXX, RAD51*/*RAD52* and histone H3.3^35^ as mentioned earlier. However, little is known about gene expression profiles for ALT+ tumors^36,37^ and as showed in our analysis, mRNA high and low expression are genomic alterations present in 76.5% of the samples tested.

Commonly, ALT is enriched in tumors of mesenchymal origin^38^, however, there is evidence of coexistence between telomerase and ALT in some solid tumors. In fact, some anti-telomerase-based treatments have shown the capability of some cells to switch to ALT and escape death^3,10^. For our study, we found the highest frequency means of genomics alterations in the following cancer types: UCS, OV, UCEC, ESCA, BLCA, ACC, SKCM, LUSC, STAD and BRCA (Figure 2A). According to literature, there is variation of ALT+ tumors in different types of cancer, for instance, for UCS, Lee *et al*. 2011 reported 40% ALT+ cell lines^39^, Heaphy *et al*. 2011 reported 7% and Lee *et al*. 2018 reported 0% ^40^. In average, the following frequencies of ALT positive cell lines have been reported for the cancer types we found to have the highest frequencies of genomic alterations: OV 10% ^40,41^, UCEC 7% ^41^, ESCA 5% ^40,41^, BLCA 5% ^40,41^, ACC 18% ^34,41^, SKCM 15% ^34,40–42^, LUSC 4% ^34,40,41^, STAD 15% ^34,40,43^ and BRCA 3% ^34,40,41^. Nonetheless, due to the difficulty and lack of sensible diagnostic techniques for ALT+ tumors, those numbers may increase ^7,44,45^.

In addition, we identified which TM genes were the most altered among the 31 PCA studies (Figure 2B) and in the top altered cancer studies (Figure 2C). This analysis, linked with the oncoprint showed in Figure 2D, correlates each altered gene with its main genomic alteration. As a result, *TP53* is the most altered gene in the majority of cases, which is not odd, due to the evidence of its function as a tumor suppressor. However, it has been reported to be co-mutated with *ATRX* and histone H3.3 ^46^ and is related with the high expression of *TERT* (a well-known factor for ALT progression) when truncated ^47^.

Along with *TP53*, we identified and proposed in this study TM genes as possible molecular targets which have been reported by Lovejoy *et al*. 2012 to be potential elements of the ALT pathway like *UBR5, NSMCE2, RFC4, MCPH1, RECQL4, LRATD2, RAD1, RAD21, NBN, FBH1, RAD54B, ATR, SMG7* and *TERF1* ^35^. Also, according to Osterwald *et al*. 2015 and Chung *et al*. 2011, *SENP5, NSMCE2, NBN, ATR* and TERF1 are involved in APBs formation^48,49^, which are common in the phenotype of ALT+ cells for replication and extension of telomere ends^50^. Identically, Dejardin *et al*. 2009, found *NSMCE2, RFC4, CCT5, NBN* and *TERF1*, to be expressed in the telomeres of ALT+ cell lines^51^.

Some of the genes proposed have been proven to play an important role in another TM mechanisms which can be ligated to ALT+ tumors, for instance: *NSMCE2* recruitment is essential for APBs functioning^48^, it is also part of the *SMC5/SMC6* complex, which its inhibition is known to disrupt APBs formation^52^. *NBN* is part of the MRN complex (*MRE11/RAD50/NBN*) which generates the G-overhang in the lead telomeric strand^53^; additionally, it promotes the ALT mechanism by recruiting *ATM* to the telomeres, allowing the invasion of adjacent telomeric DNA to be used as a template for telomere extension^54^.

Moreover, *MCPH1* and *ARID1A* bind to the telomerase reverse transcriptase (hTERT) and regulate its expression^55^, the oncoprint in Figure 2D shows these genes to be down regulated or fused across the different cancer types, which can give an insight on their role in the switching from telomerase to ALT. Other genes like *LRATD2* is known to be upregulated in cells with short telomeres^56^. *ATR* is important in the assembly of the telomerase complex^57^. *TERF1* is overexpressed in cells with long telomeres^58^ and last but not least, *RECQL4* is associated with *TERF1* in the formation of the shelterin complex^59^.

According to the TelNet database, based on their role in the ALT mechanism, TM genes are qualified as enhancers, repressors or ambiguous ^14^. Only *NSMCE2, RFC4, NBN* and *ATR* are recognized as enhancers, while, *SENP5, UBR5, RAD21, MCPH1, RECQL4, ARID1A, CCT5, LRATD2, RAD1, FBH1, RAD54B, SMG7* and *TERF1* are qualified as ambiguous. This highlights the importance of an expression analysis of the ALT-associated genes identified in this research, in order to study their role in the switching, progression or maintenance of ALT.

In order to understand the way TM proteins interact and behave in ALT+ cells we performed a protein-protein interaction network using STRING (Figure 3A). We selected and observed the most significant pathways (p< 0.001) associated with the selected TM proteins, as expected, 35% of the 103 proteins analyzed were involved in telomere organization, maintenance and telomeric DNA binding, while 22% are crucial for HDR and HRR, which altogether with non-homologous end joining (NHEJ) have an important role in the mechanism by which ALT+ cells extend its telomeres^60,61^.

Moreover, we performed an enrichment analysis for the 103 TM proteins using g:Profiler, which searches a collection of proteins with pathways, networks, gene ontology (GO) and cancer phenotypes^62^. The GO for molecular functions were DNA binding and telomeric DNA binding, the GO for biological process were telomere maintenance, DNA repair, telomere and chromosome organization. The most significant KEGG signaling pathway was HR, and the most significant REACTOME most significant pathways were DDSB repair and HRR. Finally, the human phenotype related to the proteins analyzed was abnormality of chromosome stability. As expected, the GO analysis and pathways observed were highly related to ALT progression and maintenance mechanisms.

Furthermore, we selected the proposed TM proteins in our study and the most significant pathways from the PPi and enrichment analysis and constructed a CIRCOS plot (Figure 3C) to observe the main pathways in which our candidate proteins are directly involved. As anticipated, most of the proteins like RAD54B, NSMCE2, TERF1, SMG7, RECQL4, ATR, PRKDC, RAD1, RFC4 and NBN are directly involved in pathways like DDSB, HR, HDR, HRR among others, which are crucial for the ALT mechanism.

Finally, following the procedure of López-Cortés *et al*. 2020, we used the Open Targets Platform (https://www.targetvalidation.org/)^27,63^ which allows the visualization of potential drugs targets associated with different cancer types and gives an insight about clinical trials, phase, type of drug and target, associated to each drug^63^. We found that only 9 of the TM genes we studied are included in drug trials (*ATR, PRKDC, PARP1, TOP1, SRC, POLD2, SEM1, DNMT1* and *POLE*), which highlights the need of doing more studies like these to identify new potential molecular markers to be used for ALT+ tumors diagnosis and treatment.

## Conclusions

10-15% of tumors use ALT as a TMM for telomere extension, however, the co-existence of ALT with telomerase expression has been observed in some tumors. As a consequence of anti-telomerase-based therapies in cancer treatments, cancer cells have the ability to switch from a telomerase-dependent TMM to ALT and due to the poor prognosis that ALT+ cells represent, it is very important to find a proper diagnosis and a therapeutic strategy for these cases. Nevertheless, the mechanism and factors involved in the ALT pathway are still elusive and poorly understood. In this study, we conducted a genomic analysis of 411 TM genes in 31 PCA studies and with the aid of different *in-silico* approaches, we proposed 20 candidate genes which are highly mutated and up and down regulated in different cancer types, which can be useful for future studies as possible molecular targets for the detection and understanding of the ALT pathway. Finally, we showed the need of a deep study of expression profiles and *ex vivo* assays of ALT related genes that can elucidate its role in cancer cells TMM and can provide guidance in the development of new drugs for the treatment of ALT+ tumors. We strongly believe this new decade would be promising for the study and understanding of this mechanism.

## Methods

### Gene sets

TelNet database (http://www.cancertelsys.org/telnet/) was downloaded and filtered manually to scan for genes related to the ALT mechanism. The criteria used for filtering were: TM function, significance, phenotype and annotation. *ATRX* and *DAXX* genes, already known to be involved in ALT progression were excluded from the study, resulting in 411 TM genes selected for further analysis.

### Genomic alterations

Genomic alterations (CNV amplification, CNV deep deletion, inframe mutation, truncating mutation, missense mutation, fusions, mRNA high and mRNA low) were analyzed in the cBio Portal (http://www.cbioportal.org/). A total of 9,282 samples were selected from 31 TCGA Pan-Cancer studies (LAML, ACC, BLCA, LGG, BRCA, CESC, CHOL, COAD, DLBC, ESCA, GBM, HNSC, KIRC, KIRP, LIHC, LUAD, LUSC, MESO, OV, PAAD, PCPG, PRAD, SARC, SKCM, STAD, TGCT, THYM, THCA, UCS, UCEC and UVM). The frequency means of genomic alterations were compared with a Bonferroni correction test (p< 0.05) by using SPSS Statistics Software (IBM).

### Protein-protein interaction network

In order to predict the interactions among the TM proteins we used the STRING database, with an interaction score of 0.9 (highest confidence). Most significant signaling pathways (p< 0.001) related to TMM and the ALT pathway were selected and differentiated by colors in the network.

### Gene set enrichment analysis

The set of genes was analyzed in the g:Profiler (https://biit.cs.ut.ee/gprofiler/gost), the significance threshold selected was Benjamini-Hochberg FDR (p< 0.001) and data sources consulted were GO: Molecular function, GO:Biological process, KEGG, Reactome and Human Phenotype Ontology.

## Author contribution

IAC conceived the subject, conceptualize the study and wrote the manuscript. ALC gave very valuable conceptual and scientific advices. SG and CPYM supervised the study. ALC, JGC, PGR, PEL, APV, VY, AKZ and SG did data curation and writing edition. All the authors reviewed and approved the manuscript.

## Competing interests

The authors declare no competing interests.

## Data availability statement

All data generated or analyzed for this study are included in the manuscript and its supplementary material.

## Notes

### Competing Interest Statement

The authors have declared no competing interest.

